# Synthetic Peptide Nucleic Acids in Action: *In Vivo* Interference of the *groEL* Gene from Pea Aphid Endosymbiont *Buchnera aphidicola*

**DOI:** 10.1101/2023.10.26.560802

**Authors:** Kathrine Xin Yee Tan, Shuji Shigenobu

## Abstract

The unculturable nature of intracellular obligate symbionts presents a significant challenge for elucidating gene functionality, necessitating the development of gene manipulation techniques. One of the best-studied obligate symbioses is that between aphids and the bacterial endosymbiont *Buchnera aphidicola.* Given the extensive genome reduction observed in *Buchnera*, the remaining genes are crucial for understanding the host-symbiont relationship, but a lack of tools for manipulating gene function in the endosymbiont has significantly impeded the exploration of the molecular mechanisms underlying this mutualism. In this study, we introduced a novel gene manipulation technique employing synthetic single-stranded peptide nucleic acids (PNAs). We targeted the critical *Buchnera groEL* using specially designed antisense PNAs conjugated to an arginine-rich cell-penetrating peptide (CPP). Within 24 h of PNA administration via microinjection, we observed a significant reduction in *groEL* expression and *Buchnera* cell count. Notably, the interference of *groEL* led to profound morphological malformations in *Buchnera*, indicative of impaired cellular integrity. The gene knockdown technique developed in this study, involving the microinjection of CPP-conjugated antisense PNAs, provides a potent approach for *in vivo* gene manipulation of unculturable intracellular symbionts, offering valuable insights into their biology and interactions with hosts.

## Introduction

Symbiosis refers to the establishment of mutually beneficial relationships between two or more organisms[1]. The pea aphid (*Acyrthosiphon pisum*) and its endosymbiont *Buchnera aphidicola* serve as valuable models for studying mutual symbiotic relationships and the resulting co-evolution[2]. *Buchnera* synthesizes essential amino acids to supplement the diet of the host aphid, which is composed of plant sap, whereas the host shelters the endosymbiont from harsh environments[3]. The aphid/*Buchnera* association is obligate and mutualistic, with neither partner being able to reproduce in the absence of the other. To probe gene functions and roles in this symbiosis, researchers have extensively employed molecular approaches, such as the CRISPR-Cas9 system and RNA (RNAi) interference targeting host genes[4]–[7]. In contrast to the successful genetic manipulation of the host insect, the unculturable nature of *Buchnera* poses a significant challenge, as it hinders the development of an effective gene manipulation method to study symbiont genes. This obstacle is not unique to *Buchnera* but is a common characteristic of any obligate symbiont of other symbiotic systems where symbionts cannot thrive outside their host environments. Consequently, the pursuit of effective gene manipulation strategies for obligate symbionts remains a challenge.

Peptide nucleic acids (PNAs) are synthetic analogs of DNA or RNA and have shown promise as antimicrobial therapeutic agents. Extensive research has tested the efficacy of PNAs against a range of pathogenic bacteria, including Gram-negative bacteria such as *Escherichia coli*, *Klebsiella pneumoniae,* and *Salmonella Typhimurium* and gram-positive bacteria such as *Streptococcus pyrogenes* and *Bacillus subtilis*[8], [9]. PNAs have a unique structure in which the deoxyribose phosphate backbone is replaced with a pseudopeptide backbone attached to the nucleobases. Structural modification confers several advantages that enhance the utility of PNAs as antimicrobial agents. First, they are chemically stable and resistant to degradation by nucleases and proteases[10]. Second, owing to the lack of charge repulsion, the PNA-DNA and PNA-RNA duplexes are more stable than natural DNA-DNA or RNA-RNA duplexes. Additionally, the unique structure of PNAs facilitates high-specificity binding to complementary DNA and RNA sequences. This enables meticulously designed antisense PNAs to effectively interfere with the expression of target genes. For instance, Good and Nielson (1998) demonstrated the effectiveness of antisense PNAs targeting the start codon regions of *E. coli* β-galactosidase (*lacZ*) and β-lactamase (*bla*) genes, resulting in significant inhibition of gene expression[11]. Furthermore, antisense PNAs have demonstrated the capability to selectively kill targeted bacteria in mixed cultures containing multiple bacterial species[8]. Taken together, their endurance against enzymatic degradation, along with their precise gene interference capabilities and selective bacterial targeting, make PNAs a promising avenue for future drug development research.

Although PNAs have significant potential for antimicrobial drug development, a persistent challenge lies in the effective translocation of the molecule across cell boundaries. However, this obstacle has been substantially mitigated by conjugating PNAs with cell-penetrating peptides (CPPs) to facilitate their uptake across the bacterial cell wall[8], [9], [12], [13]. CPPs are short peptides rich in basic amino acids such as lysine and arginine[14]. CPPs have been widely used to deliver large cargo molecules, including oligonucleotides, across the cell barrier without inducing substantial membrane damage. Therefore, selecting a suitable CPP conjugate for antisense PNAs is crucial for ensuring the effective internalization of PNAs into targeted cells. This approach ensures efficient delivery of antisense PNAs and enhances their potential for gene interference applications.

In the present study, we developed a novel protocol for *in vivo* gene interference in unculturable intracellular symbionts using antisense PNAs. Our target gene was *Buchnera groEL*, which encodes the heat shock protein GroEL, also known as symbionin. This gene was selected because of its extremely high expression in *Buchnera*[15], [16]. GroEL is hypothesized to play a crucial role as a molecular chaperone in assisting the folding of symbiont proteins[16], [17]. Therefore, we anticipated that interference with the *groEL* gene would have a significant effect on the endosymbiont. We carefully designed antisense PNA and selected a compatible CPP to facilitate specific knockdown and efficient delivery. Microinjection of CPP-conjugated antisense PNAs into the host insect was performed, and subsequent molecular and microscopic analyses showed successful downregulation of *groEL* expression, accompanied by a decrease in *Buchnera* cells and observable morphological changes in the endosymbiont. To the best of our knowledge, this is the first instance of gene knockdown in obligate symbionts, offering a pioneering approach for the study of unculturable symbionts.

## Results

### Design of peptide conjugated antisense PNAs

We designed antisense PNAs to be complementary to the region encompassing −5 to +5 bases from the translational start codon (ATG) of the *Buchnera groEL* gene (Figure 1). It is well known that PNAs targeting the translational start codon and the Shine-Dalgarno region demonstrate higher efficiency compared to antisense PNAs targeting other gene sequence regions [18]. We designed the PNAs sequence in 10 bases, considering the balance between sequence specificity and constraints of cellular uptake[19]. A commonly employed strategy to facilitate cellular penetration of antisense PNAs involves conjugation with CPP. In particular, arginine-rich CPPs are favored because of their low toxicity and high effectiveness in PNA delivery[20]. Hence, we appended an arginine-rich CPP, (RXR)_4_XB, to the N-terminus of the PNAs (Figure 1, Table 1).

**Figure 1:**
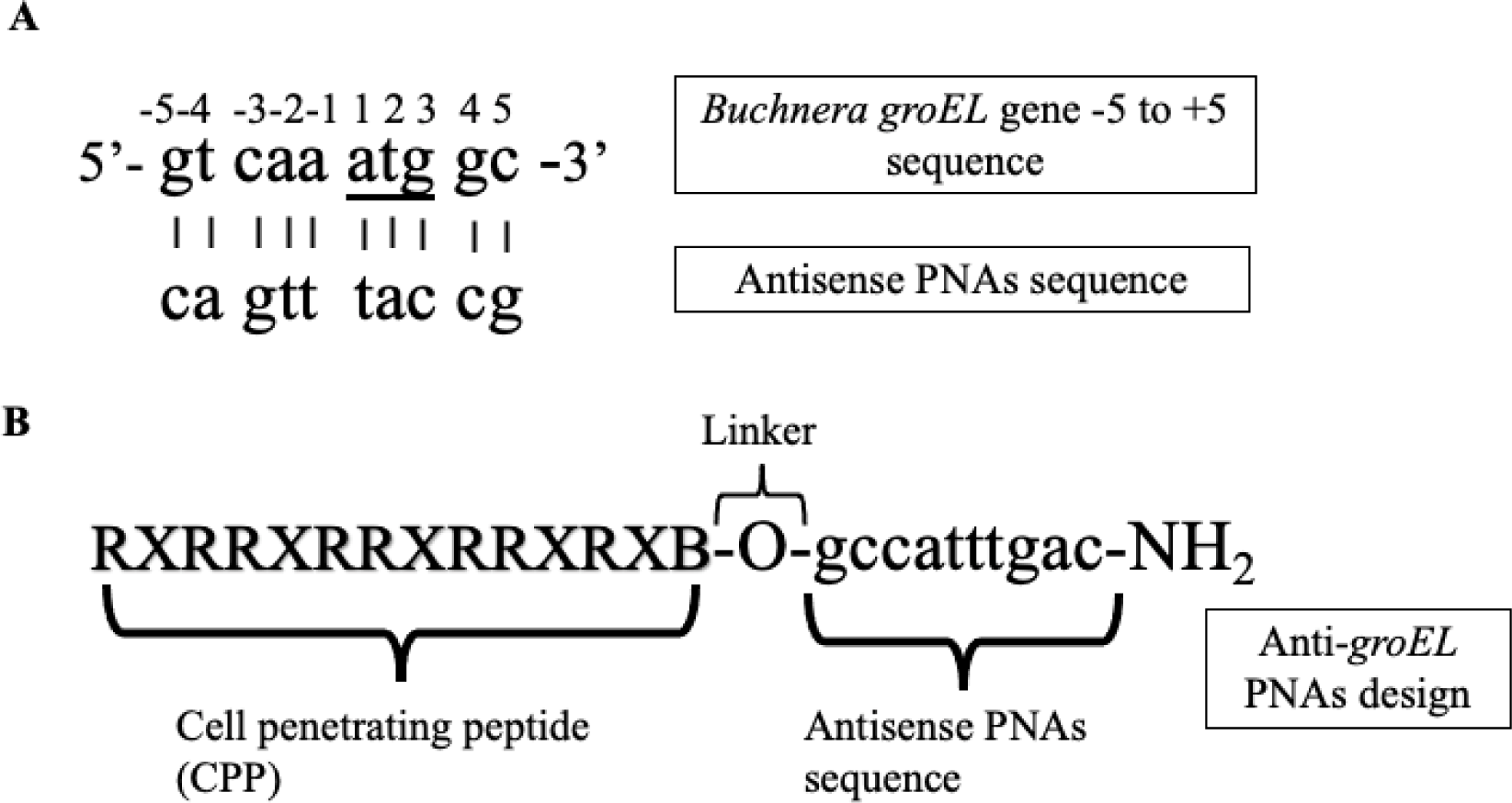
Design of peptide-conjugated antisense PNAs for *Buchnera groEL*. The intervention site within *Buchnera groEL*, ranging from position −5 to +5, was selected for the synthesis of peptide-conjugated antisense PNAs. Panel (A) depicts the *Buchnera groEL* sequence from −5 to +5, including the reverse complementary PNAs sequence; the start codon, ATG, is underlined. The design of anti-*groEL* PNAs incorporates an arginine-rich cell-penetrating peptide (CPP) conjugated to the antisense PNAs sequence (B).

**Table 1:**
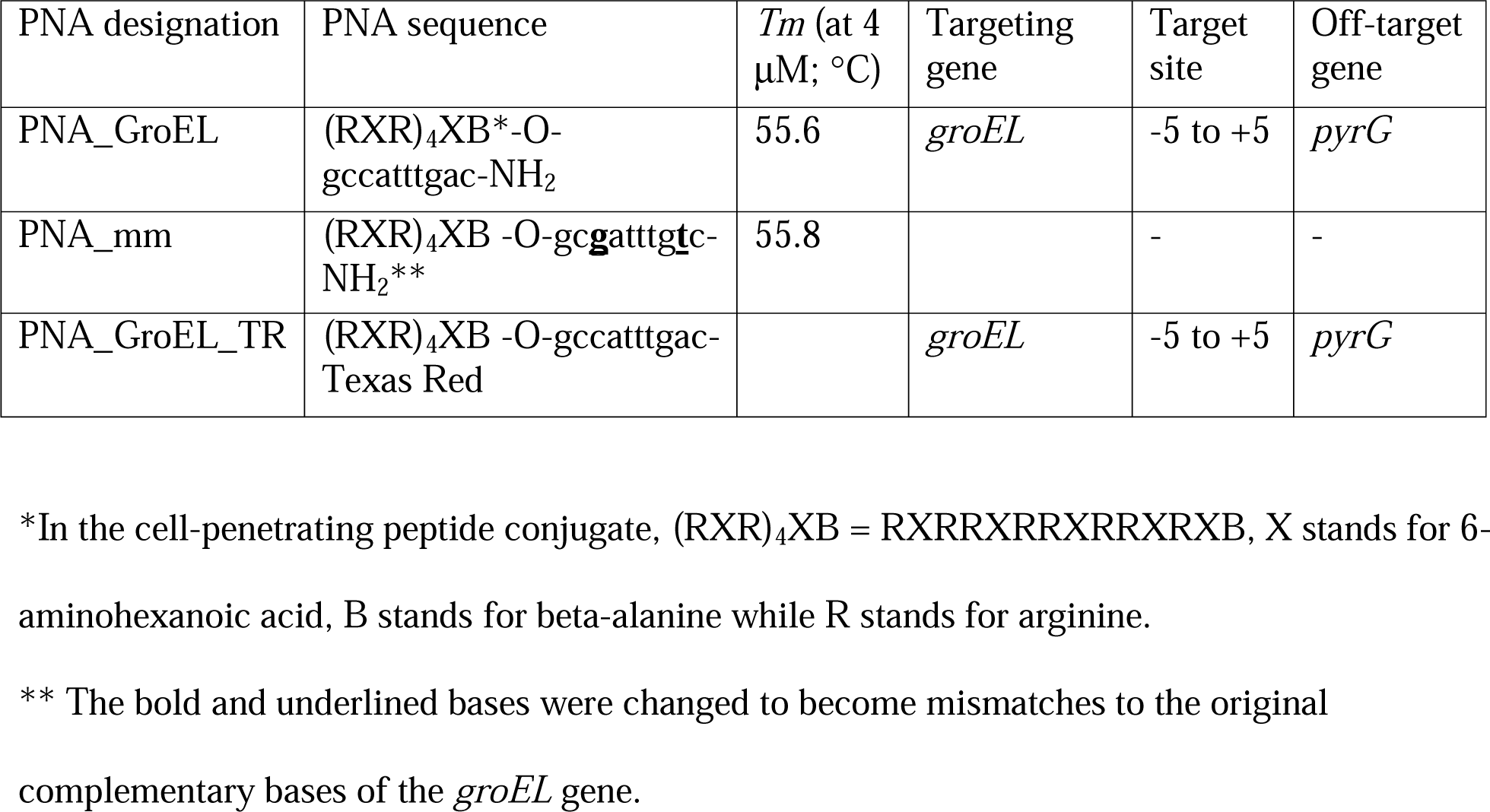
*Buchnera* genes targeting PNAs.

| PNA designation | PNA sequence | *T<sub>m</sub>* (at 4<br>μM; °C) | Targeting<br>gene | Target<br>site | Off-target<br>gene |
| --- | --- | --- | --- | --- | --- |
| PNA_GroEL | (RXR)<sub>4</sub>XB*-O-gccatttgac-NH<sub>2</sub> | 55.6 | *groEL* | -5 to +5 | *pyrG* |
| PNA_mm | (RXR)<sub>4</sub>XB -O-gc**g**atttg<u>t</u>c-NH<sub>2</sub>** | 55.8 |  | - | - |
| PNA_GroEL_TR | (RXR)<sub>4</sub>XB -O-gccatttgac-Texas Red |  | *groEL* | -5 to +5 | *pyrG* |

\*In the cell-penetrating peptide conjugate, (RXR)<sub>4</sub>XB = RXRRXRRXRRXRXB, X stands for 6-aminohexanoic acid, B stands for beta-alanine while R stands for arginine.
\*\* The bold and underlined bases were changed to become mismatches to the original complementary bases of the *groEL* gene.

### Peptide-conjugated anti-*groEL* PNAs reduced *Buchnera* titer in the pea aphid

Given their massive expression and conservation across the primary endosymbionts of a wide variety of insect lineages, GroEL homologs have been postulated to be essential genes for endosymbionts, including *Buchnera*[21]. First, we investigated the effect of anti-*groEL* PNAs (PNA_GroEL) on *Buchnera* survival. We employed quantitative PCR (qPCR) to evaluate the *Buchnera* titer in aphid nymphs subjected to treatment with PNA_GroEL and PNAs with mismatches (PNA_mm). To ensure accurate quantification, we used the aphid single-copy gene *rpL7* as a reference to normalize the *Buchnera* single-copy gene *dnaK*. PNA_GroEL represents CPP-conjugated anti-*groEL* PNAs, whereas PNA_mm served as the corresponding negative control, featuring the same construct, but with two mismatched bases in the PNAs sequence (Figure 2). *Buchnera* titer was quantified 24 h after injection. A significant reduction was observed in the *Buchnera* titer in aphid nymphs treated with PNA_GroEL (*M* = 1.43, *SD* = 0.173) compared to those treated with PNA_mm (*M* = 1.55, *SD* = 0.141) (*t* (43) = −2.484, *p* = 0.008483), indicating a noteworthy impact of anti-*groEL* PNA on *Buchnera* survival or growth.

**Figure 2:**
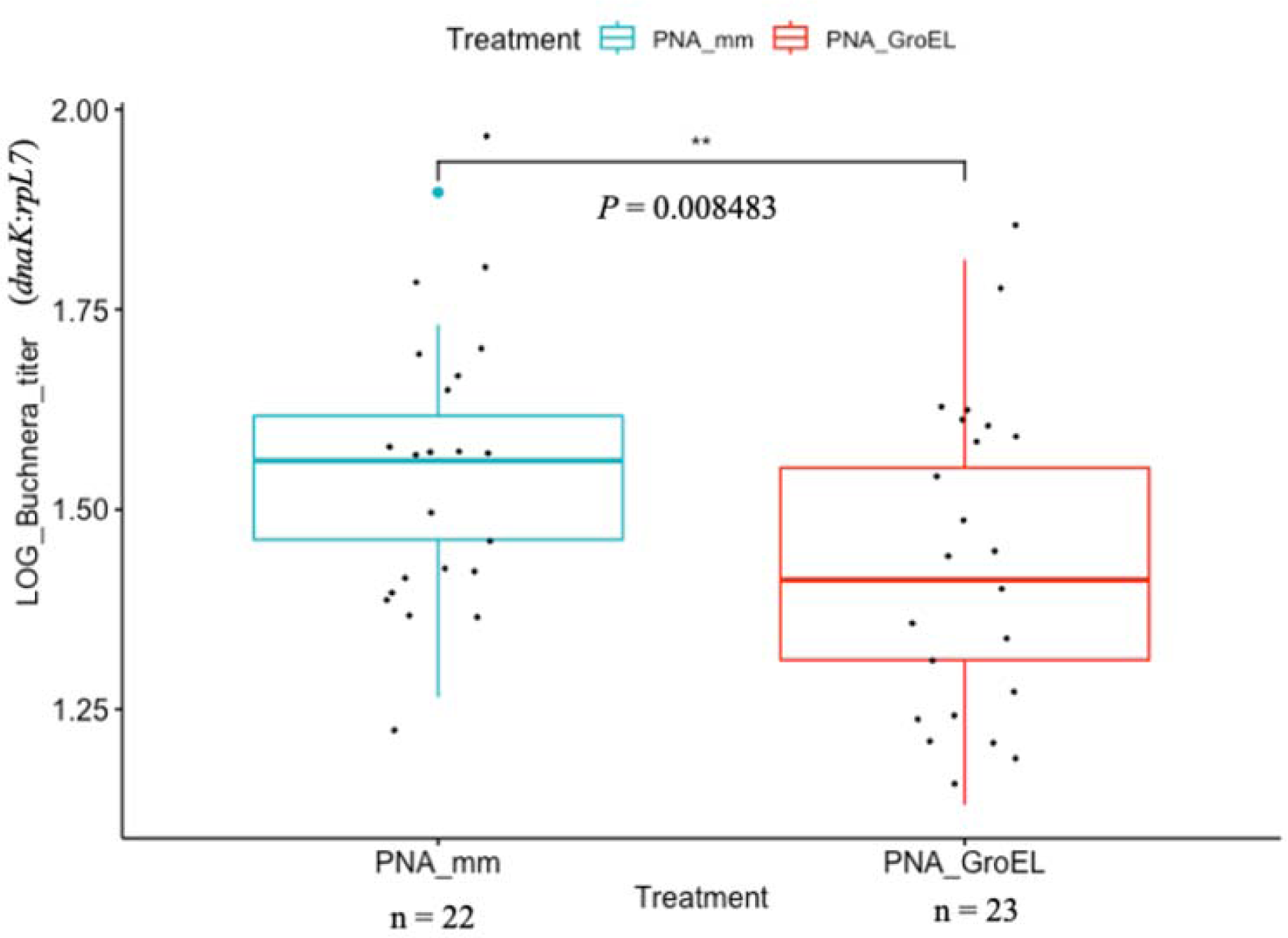
Comparative analysis of *Buchnera* titer in aphid nymphs treated with PNA_GroEL and PNA_mm. *Buchnera* titer in aphid nymphs treated with PNA_GroEL is significantly lower compared to those treated with PNA_mm. Second instar aphid nymphs were injected with 15 μM PNA_GroEL (red boxplot) or control PNA_mm (blue boxplot). *Buchnera* titer was quantified using qPCR 24 h after PNA injection. The data were log-transformed and tested with a one-tailed Student’s t-test. *P* value is shown between the two boxplots (*p =* 0.008483 < 0.01). Each black dot indicates a single PNA-treated aphid nymph.

### Peptide-conjugated anti-*groEL* PNAs reduced *Buchnera groEL* expression

The abundance of *groEL* transcripts was expected to decrease due to the inhibitory effect of complementary antisense PNAs. To verify this, we compared the transcript levels of *Buchnera groEL* between PNA_GroEL-and PNA_ mm-treated aphid nymphs using reverse transcription-quantitative polymerase chain reaction (RT-qPCR) (Figure 3). Upon administration of ten μM PNA_GroEL, we observed a significantly lower expression of the *Buchnera groEL* gene in aphid nymphs (*M* = 21.2, *SD* = 9.34), compared to those treated with the same concentration of PNA_mm (*M* = 28.8, *SD* = 8.90) (*t* (50) = −2.9897, *p* = 0.002162), thus confirming the knockdown effect of anti-*groEL* PNAs on *Buchnera groEL* expression.

**Figure 3:**
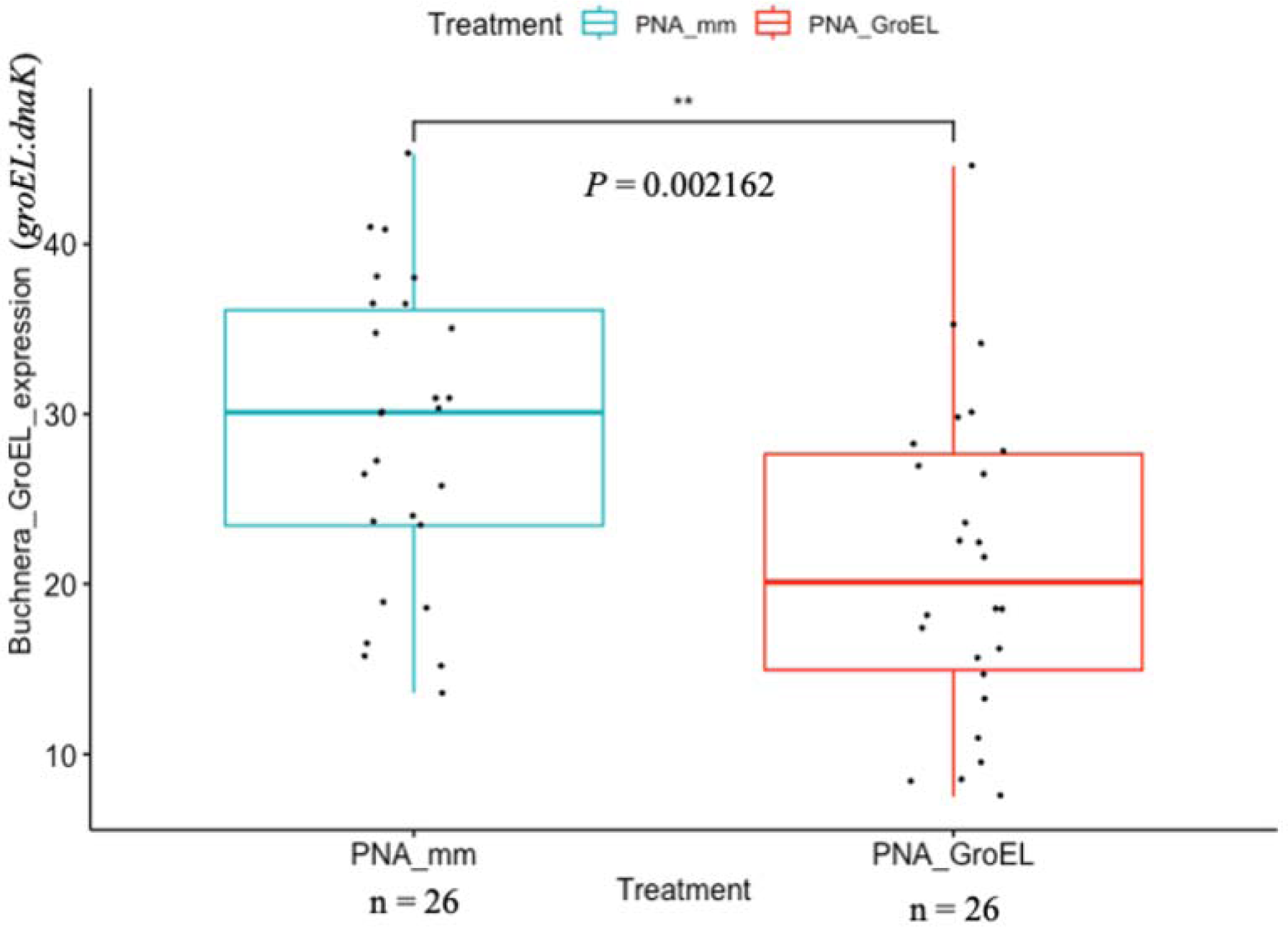
Differential *Buchnera groEL* gene expression in aphid nymphs treated with PNA_GroEL and PNA_mm. *Buchnera groEL* gene expression in aphid nymphs treated with PNA_GroEL was significantly lower than that in those treated with PNA_mm. Second instar aphid nymphs were injected with 10 μM peptide-conjugated antisense *groEL* PNA (PNA_GroEL; red boxplot) or control PNAs (PNA_mm; blue boxplot). The *groEL* gene expression was quantified using RT-qPCR 24 h after PNA injection. Statistical significance of *Buchnera groEL* gene expression was tested with a one-tailed Student’s t-test (*p* = 0.002162 < 0.01). Each black dot indicates a single PNA-treated aphid nymph.

### Delivery of peptide-conjugated anti-*groEL* PNAs into bacteriocytes and *Buchnera* cells

Effective delivery of PNAs across cellular membranes poses a significant challenge, especially because *Buchnera* resides within a specialized host cell, the bacteriocyte. To trace the uptake and transport of the peptide-conjugated anti-*groEL*, we used PNAs conjugated with the fluorescent dye Texas Red. Texas Red signal was detected in the bacteriocytes of aphid nymphs treated with PNA_GroEL_TR 24 h after injection (Figure 4D-F). High-resolution images obtained by super-resolution microscopy distinctly exhibited dispersed signals of Texas Red within the aphid bacteriocytes and *Buchnera* cells (Figure 4F’–F”). The fluorescent signal was also detected within the bacteriocytes of the treated nymphs as early as approximately six hours following injection (see Supplementary Figure S1). Notably, Texas Red signals were not confined to bacteriocytes but were also discernible in other aphid tissues, such as fat body cells, gut cells, and even within embryos developing inside parthenogenetic ovarioles (see Supplementary Figure S2). This collective evidence supports the successful delivery and translocation of peptide-conjugated PNAs into bacteriocytes, and subsequently into endosymbionts.

**Figure 4:**
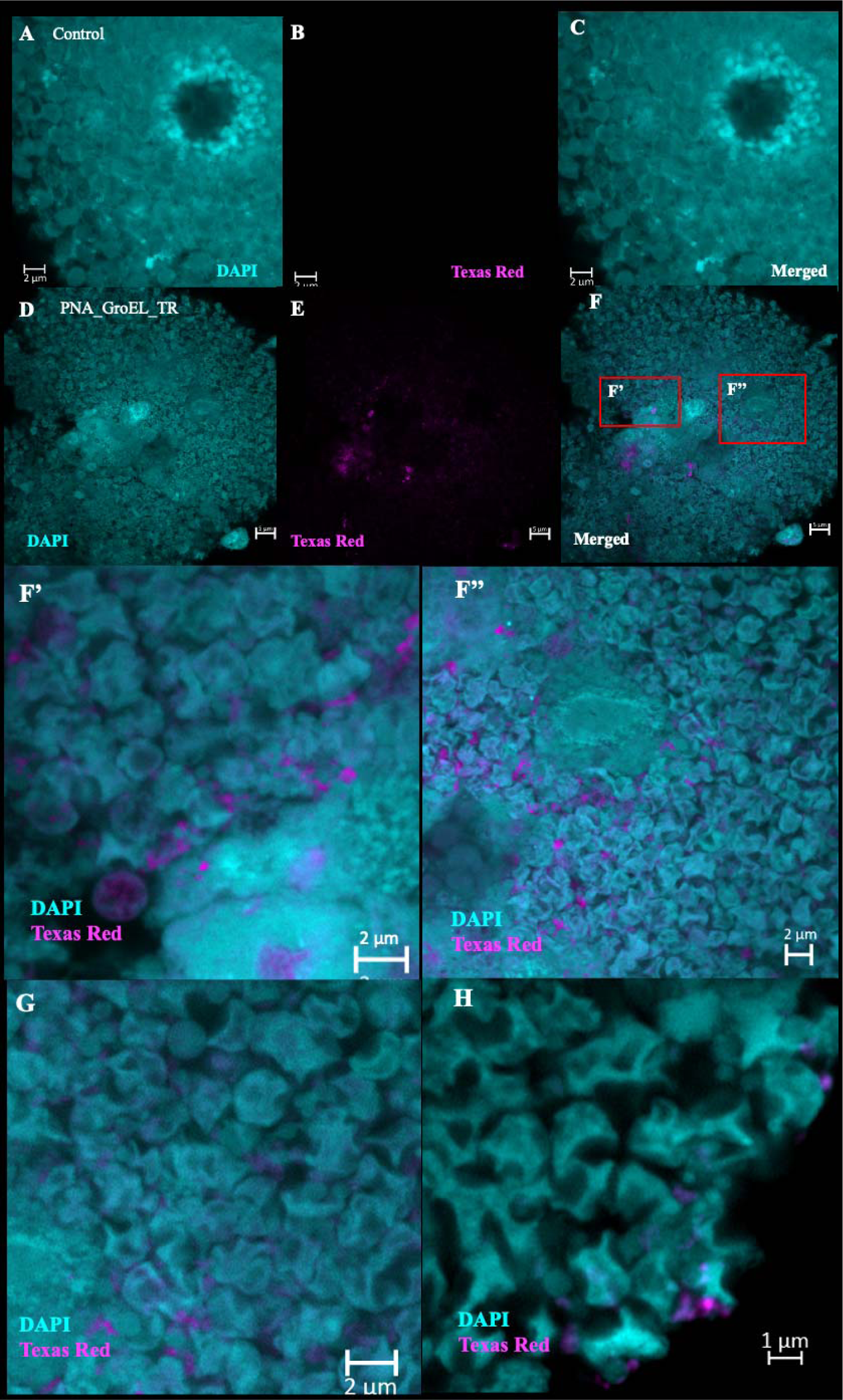
Super-resolution imaging of aphid bacteriocytes and *Buchnera* cells reveals PNA_GroEL_TR effects. Aphid nymphs were injected with CaCl_2_ solution (12 mM) or PNA_GroEL_TR (10 μM in 12 mM CaCl_2_ solution). Injected aphid nymphs were dissected after 24 h. Dissected bacteriocytes were fixed and stained with a DAPI solution to mark the nuclei of bacteriocytes and the chromosome of *Buchnera*. Aphid nymphs injected with CaCl_2_ solution injected were used as the control (A-C). Detection of the Texas Red fluorescent signal in bacteriocytes and *Buchnera* cells indicated the successful penetration of PNAs into the cells (D-F). Zoom-in images showed that the Texas Red signal was scattered in the bacteriocytes (F’-F’’). *Buchnera* cells were distorted in the PNA_GroEL_TR treated sample (G-H).

### *Buchnera* cell malformation was observed in aphid nymphs treated with peptide conjugated anti-*groEL* PNAs

*Buchnera* cells presented a spherical and round morphology (Figure 4A-C) under normal conditions[22]. However, our microscopic investigation of the PNA_GroEL_ TR-treated samples revealed severe malformation of *Buchnera* cells (Figure 4G-H). These *Buchnera* cells appeared wrinkled and deviated from their typical spherical form, with some exhibiting a crescent shape.

To quantify the extent of *Buchnera* cell deformation from the imaging data, we used the SymbiQuant software. This machine learning-based object detection tool was designed for the identification and enumeration of endosymbionts in host tissue micrographs[23]. The software authors trained SymbiQuant to detect *Buchnera* cells using microscopic images captured during normal aphid development[23]. Therefore, we surmised that SymbiQuant primarily identifies and counts spherical *Buchnera* cells, which is the morphology of normal cells, consequently distinguishing these from those exhibiting abnormal morphologies. As anticipated, Figures 5B and 5C (examples of output images after SymbiQuant processing) show that SymbiQuant failed to recognize malformed *Buchnera* cells but detected normally shaped cells with high accuracy.

**Figure 5:**
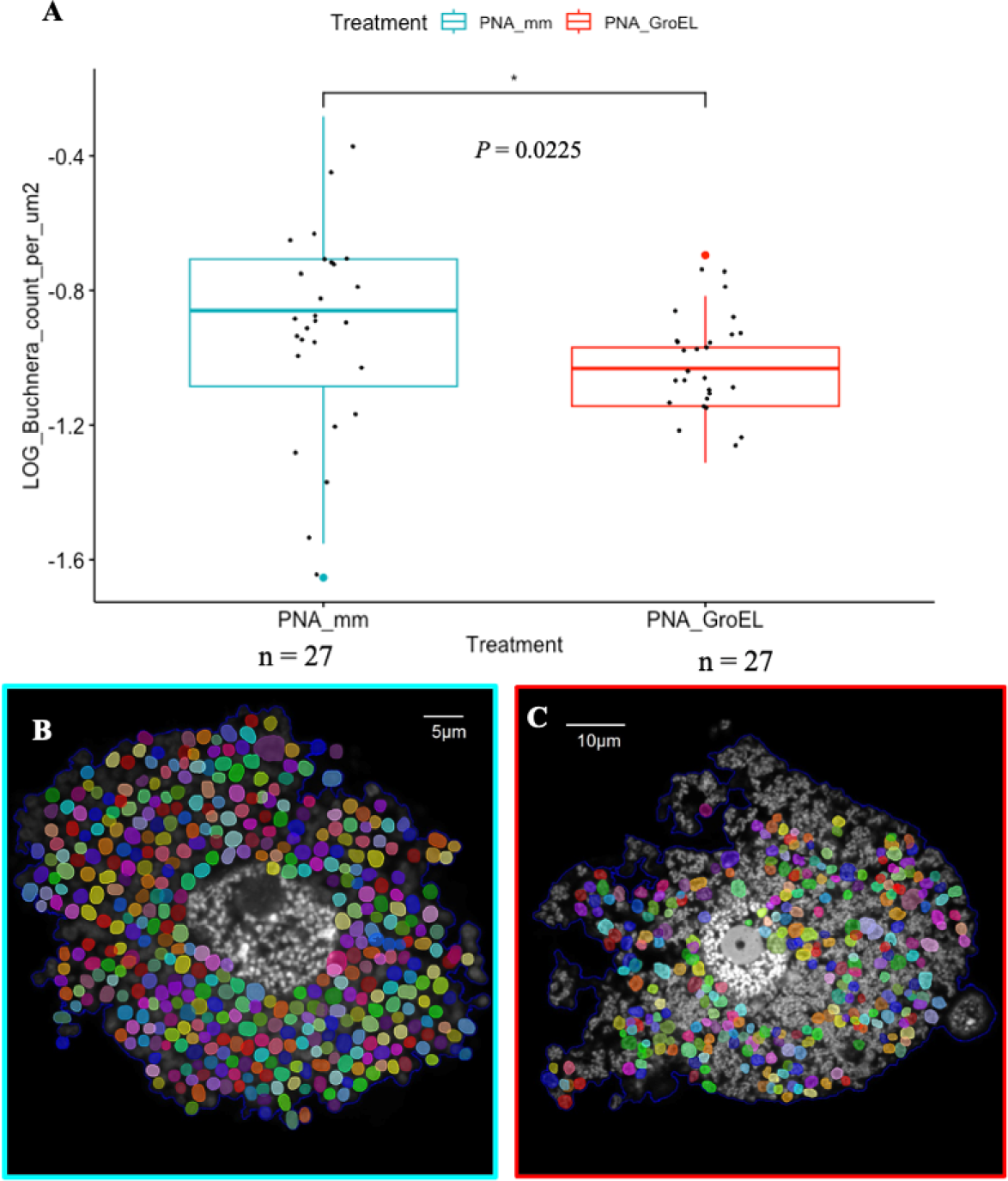
Quantification of *Buchnera* cells using machine-learning tool, SymbiQuant. The log-transformed *Buchnera* count per μm^2^ cell area was significantly lower in aphid nymphs injected with PNA_GroEL than PNA_mm (*p* = 0.0225 < 0.05, one-tailed Welch Student’s t-test) (A). *Buchnera* cells successfully detected by SymbiQuant were marked by multi-coloured circles (B: PNA_mm; C: PNA_GroEL). Injected aphid nymphs were dissected after 24 h. Dissected bacteriocytes were fixed and stained with a DAPI solution to mark the nuclei of bacteriocytes and the chromosome of *Buchnera*. Single optical plane 1024 x 1024 pixels confocal images were taken with a 100x oil immersion lens. *Buchnera* count detected using SymbiQuant was normalised by cell area measured by ImageJ (*Buchnera* count per μm^2^) followed by log transforming the data. Each black dot represents the data from a single image.

We compared *Buchnera* counts within bacteriocytes between aphid nymphs treated with PNA_GroEL and PNA_mm (control) via SymbiQuant (Figure 5). Using the pretrained SymbiQuant model, to detect the spherical-*Buchnera* cells, we observed a significantly lower *Buchnera* density in the PNA_GroEL-treated samples (*M* = −1.04, *SD* = 0.139) than in the control group treated with PNA_mm (*M* = −0.908, *SD* = 0.312) (*t* (35.947) = −2.0771, *p* = 0.0225). These observations suggested that anti-*groEL* PNA treatment significantly reduced the population of *Buchnera* cells with normal morphology, promoting the prevalence of cells with abnormal morphologies. This could indicate metabolic and/or cellular abnormalities in *Buchnera* as a result of PNAGroEL treatment.

### Evaluating the off-target effect of anti-*groEL* PNAs on untargeted genes

A critical aspect of the PNA design involves ensuring the uniqueness of the target site. However, owing to the short length of PNAs, full or partial complementary matches with genome regions other than the target gene occur frequently[24]. Our comprehensive genome-wide search revealed that the 10-bp anti-*groEL* PNA exhibited no sequence matching with any *Buchnera* genes, except the target *groEL*, at their translational start codons or Shine-Dalgarno regions. Nevertheless, the sequence of the anti-*groEL* PNA did match a middle segment of the *pyrG* gene. To address potential off-target effects, we evaluated whether PNAs designed to target the *groEL* gene also inhibited the expression of the *pyrG* gene. RT-qPCR was performed to assess the expression of the *pyrG* gene (Figure 6A). Our results showed that the expression of the *pyrG* gene in PNA_GroEL (*M* = 0.429, *SD* = 0.378) and PNA_mm-treated (*M* = 0.688, *SD* = 0.541) aphid nymphs was not significantly different (Mann–Whitney U = 17, n_1_ = n_2_ = 7, *p =* 0.3829).

**Figure 6:**
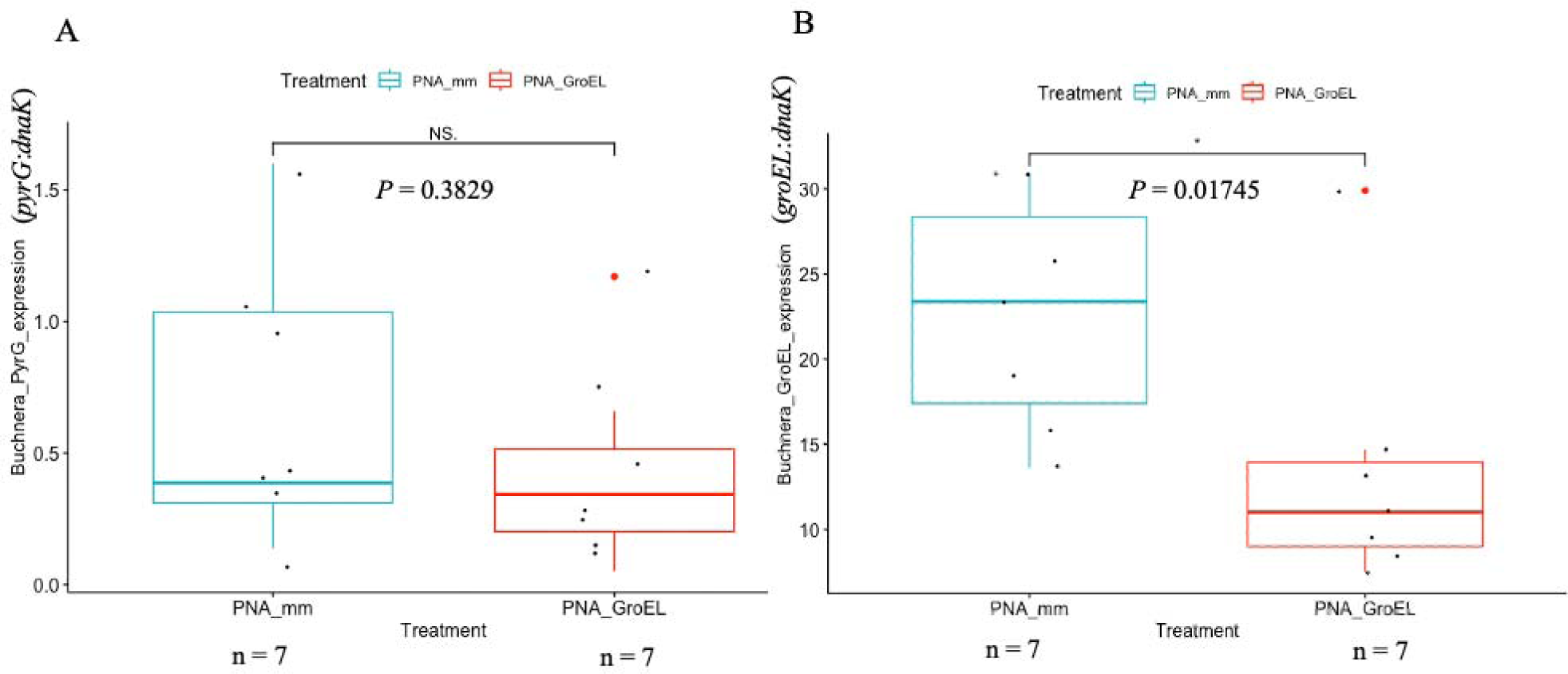
Comparative analysis of *Buchnera pyrG* expression in PNAs-injected aphid nymphs. Second instar aphid nymphs were injected with 10 μM peptide-conjugated antisense *groEL* PNA (PNA_GroEL; red boxplot) or control PNAs (PNA_mm; blue boxplot). *Buchnera pyrG* gene expression was quantified using RT-qPCR 24 h after PNAs injection. A: The difference in *Buchnera pyrG* gene expression between aphid nymphs treated with PNA_GroEL and PNA_mm is not significant (*p* = 0.3829 > 0.05, two-tailed Mann-Whitney U test). B: *Buchnera groEL* gene expression was significantly lower in the PNA_GroEL treated group in the same RNA samples (*p* = 0.01745 < 0.05, one-tailed Student’s t-test). Each black dot indicates a single PNAs treated aphid nymph.

Concurrently, the expression of the *groEL* gene was significantly lower in the PNA_GroEL group (*M* = 13.5, *SD* = 7.68) than in the PNA_mm group (*M* = 22.8, *SD* = 6.94) when the same RNA samples were analyzed (*t* (12) = −2.3777, *p* = 0.01745) (Figure 6B). These data suggest that the anti-*groEL* PNA selectively targets and inhibits *groEL* expression without significantly affecting the expression of the *pyrG* gene, hence demonstrating a high degree of specificity.

## Discussion

The absence of an effective genetic tool to manipulate gene expression in endosymbionts has significantly hindered our understanding of the functions of genes that persist in reduced genomes, particularly given that these symbionts are typically unculturable outside their host organisms. This has impeded our understanding of the molecular mechanisms that govern mutualistic relationships. In this study, we focused on *Buchnera,* an aphid endosymbiont, as a representative model of unculturable obligate symbionts. We formulated a novel *in vivo* protocol for gene interference in this endosymbiont using antisense PNAs. We primarily focused on the *groEL* gene in *Buchnera*, which encodes the heat shock protein GroEL (or symbionin). Since this protein is known to play a crucial role in symbiosis with host insects, we anticipated that interference would lead to discernible phenotypic changes in the symbiont. We designed an antisense PNA targeting the translational initiation site to facilitate specific knockdown, and opted for a suitable CPP for efficient delivery. CPP-conjugated antisense PNAs were microinjected into host insects. Molecular and microscopic evaluations revealed marked downregulation of *groEL* expression, which was associated with a reduction in *Buchnera* cell numbers and evident morphological malformations in the endosymbiont. Additionally, we assessed the off-target effects to validate the specificity of our PNA design. To the best of our knowledge, this is the first report of a successful gene knockdown in obligate symbionts.

To maximize the utility of PNAs as antimicrobial agents, efforts have been made to enhance their delivery efficiency and specificity. One important factor to consider is the length of PNAs, which can affect their uptake and effectiveness. The optimum length of PNAs for effective uptake by *Escherichia coli* is 10 to 12-mers[19]. The increase in PNAs’ length corresponded to the uptake rate of PNAs, as evidenced by the reduction in *E. coli* inhibition. The target site of our anti-*groEL* PNAs was deliberately selected at the −5 to +5 region of the translational start site of the gene. This region was selected because the start codon and Shine-Dalgarno motif are known to be particularly sensitive to antisense PNA treatments. However, PNAs targeting regions outside of this specific region are generally futile[18]. Therefore, we suggested that blocking the translational initiation site halts ribosome assembly making the site more sensitive to treatment with antisense agents.

Another crucial factor in designing PNAs is the selection of an appropriate CPP for conjugation to the antisense oligonucleotides. This choice significantly affects the delivery efficiency of the molecule across cellular membranes. In this study, we carefully selected arginine-rich CPP from among these candidates. Some CPPs, including the widely used (KFF)_3_K-peptide, rely on specific transporters, such as SbmA, an inner membrane protein, for efficient cell penetration[25]. However, *Buchnera* has an extremely reduced genome, lacking most of the transporters conserved in enterobacteria; therefore, CPPs that require specific transporters should be avoided. Because the SbmA-encoding gene is not present in the *Buchnera* genome, we must ensure that our choice of CPP conjugate works in an SbmA-independent manner[26]. By selecting an arginine rich CPP, the (RXR)_4_XB peptide, we were able to overcome the reliance on the SbmA transporter. The (RXR)_4_XB peptide is internalized by cells via various mechanisms, including direct translocation, endocytosis, and membrane fusion. This is achieved through the interaction of the cationic (RXR)_4_XB peptide with anionic groups, such as phosphates, carboxylates, and sulfate groups, present on the cell membrane[27]. The (RXR)_4_XB peptide has been reported to function in an SbmA-independent manner and has demonstrated higher PNAs delivery efficiency and lower cell toxicity, making it an excellent choice for use in *Buchnera* and other endosymbionts that often have streamlined genomes with a reduced repertoire of transporter genes. [20], [28]. Additionally, the Kaplan-Meier survival analysis we performed showed that over 60 % of aphid nymphs treated with peptide-conjugated PNAs could survive for more than seven days, indicating a low pernicious impact of the CPP-conjugated PNAs on the host (see Supplementary Figure S5).

The use of fluorescent-tagged PNAs allowed the visualization of their penetration into the target cell. In this study, Texas Red was conjugated to the C-terminus of anti-*groEL* PNAs. We expected that CPP-conjugated PNAs would penetrate cells indiscriminately rather than being selective towards our target endosymbionts within bacteriocytes. Indeed, we detected Texas Red signals in embryos, fat cells, gut tissues, and bacteriocytes (see Supplementary Figure S2), indicating a broad non-cell-specific delivery of PNAs. Notably, PNAs were detected within the *Buchnera* cells. This indicates that the PNAs not only traversed the cellular membrane of the host bacteriocytes but also penetrated the bacterial membranes as well as the symbiosomal membranes, which are host-derived membranes that encapsulate endosymbiont cells. The bacterial membrane system consists of an outer membrane, peptidoglycan layer, and inner membrane. Thus, CPP-conjugated PNAs have the ability to penetrate both eukaryotic and bacterial membranes, ensuring delivery into endosymbionts.

*groEL* is a highly conserved gene that encodes the heat shock protein GroEL, which belongs to the HSP60 family. In *Buchnera*, its homolog, also known as symbionin, is expressed at such an exceptionally high level that it overwhelmingly dominates the entire proteome of *Buchnera* under symbiotic conditions, even in the absence of any discernible stress stimuli[15], [29]. This pronounced expression of GroEL homologs is not unique to *Buchnera* but is observed in many endosymbionts in symbiotic settings[15], [16]. Furthermore, Fares et al. (2002) reported that *Buchnera* GroEL experienced an accelerated rate of amino acid substitution and fixation by the positive selection of amino acids replaced at the peptide- and GroES-binding sites in GroEL. Consequently, it has been hypothesized that GroEL homologs in *Buchnera*, and more broadly, in endosymbionts, play pivotal roles in the symbiotic relationship with the host[16], [17]. However, to date, direct evidence for this has been elusive because of the absence of viable gene knockdown or knockout techniques for obligate symbionts. To the best of our knowledge, this study offers the first direct evidence that underscores the essential role of *groEL* in *Buchnera* by using a gene knockdown technique involving PNAs. Successful inhibition of *groEL* expression resulted in a decrease in the number of healthy *Buchnera* cells within aphids treated with CPP-conjugated anti-*groEL* PNAs. Microscopic observations revealed morphological malformations in *Buchnera* cells after *groEL* interference, which is indicative of the compromised cellular integrity of endosymbionts. While the exact mechanism remains uncertain, a plausible explanation is that given its function as a molecular chaperone, interfering with GroEL might disrupt the appropriate folding of a wide variety of proteins within *Buchnera* cells, adversely affecting their survival.

Owing to the length constraints (10-12 bp as discussed above) for effective PNAs delivery, there is a potential risk that the designed PNAs might align with untargeted genes. Therefore, we carefully examined the off-target effects. Ideally, there should be at least two base mismatches, as a single-base mismatch may still allow PNAs to bind to untargeted genes, albeit with a greatly reduced binding affinity[8], [30]. In fact, the anti-*groEL* PNAs that we devised perfectly reverse-complementary matched an untargeted gene, *pyrG* (see Supplementary Table S2). However, this alignment occurs in the middle of the coding region and does not correspond to the translational start site, which is particularly sensitive to reverse-complementary inhibition by PNAs. To address concerns regarding the impact of our anti-*groEL* PNAs on the untargeted gene *pyrG*, we performed RT-qPCR and confirmed that the expression of the *pyrG* gene was unaffected by the anti-*groEL* PNAs (Figure 6A). In addition to the *pyrG* gene, we analyzed the expression of *rrs* and *rpoA* genes, which have sequences entirely dissimilar to those of the anti-*groEL* PNAs (see Supplementary Figure S4). As expected, neither gene exhibited notable downregulation in expression in the PNA_GroEL-treated samples compared to the control samples (PNA_mm), although a significant elevation in *rrs* expression in the PNA_GroEL group seems to be peculiar.

The successful interference of the *Buchnera groEL* gene with CPP-conjugated antisense PNAs is an important milestone in exploring the functionality of genes in endosymbionts. The methodology established for *Buchnera* may be extended to gene knockdown studies in other obligate symbionts. The strategy of PNA design (e.g., targeting translation start sites) and the choice of CPP should be adaptable for use in different obligate symbiotic contexts. However, the administration of PNAs to host organisms poses a set of challenges that require customized optimization for each specific system. Despite the high post-treatment survival rate, we cannot assert that the PNAs treatment has no pernicious effect on the host, especially when the PNAs are used at a higher concentration or for a prolonged period. More studies are required to understand the subtle impacts of PNAs towards the host. Although we analysed only the *groEL* gene, theoretically, our method can be applied to any gene. In summary, the microinjection of CPP-conjugated antisense PNAs is a potential approach to undermine gene function in unculturable obligate endosymbionts, paving the way for more in-depth symbiotic research.

## Materials and methods

### Aphid rearing

In this study, we utilized a parthenogenetic clone of the pea aphid *A. pisum* str. *ApL,* which was obtained from Sapporo, Hokkaido, Japan (known as Sap05Ms2 in [31]). Viviparous aphids were maintained on broad bean plants (*Vicia faba* L.) at 16 °C with a photoperiod of 16/8 h light/dark. Four viviparous adult female aphids were randomly selected and placed in culture cups containing young broad bean plants. The adult female aphids were allowed to reproduce for 48 h before being removed from the culture cup. The offspring aphid nymphs in each culture cup were then allowed to undergo a 48-hour growth phase. The ages of the aphid nymphs used in the experiment were estimated to range from 72 to 96 h, corresponding to 3 to 4 d of age. Specifically, we selected aphid nymphs at the second instar stage (N2), encompassing the period from day 3 to day 6 after birth[32].

### Design of *Buchnera groEL* targeting peptide conjugated antisense PNAs

The general criteria for designing PNAs were detailed in the Results section. To obtain the *Buchnera groEL* gene sequence, we used *Buchnera aphidicola* str. APS [33] (GenBank accession number: NC_002528.1; 18715 bp – 20361 bp). Subsequently, peptide-conjugated anti-*groEL* PNAs (PNA_groEL) and PNAs containing mismatched bases for anti-*groEL* (referred to as PNA_mm) were designed. To ensure specificity, we used BLAST to identify potential non-specific target sites for our PNAs sequences. To facilitate the detection of nonspecific target sites and aid in the design of PNAs targeting the translational start site of a gene, we developed a customized Python script. Additionally, for user convenience, we prepared a Google Colab notebook containing a Python script (see the Supplementary file). Synthesis and purification of the PNAs were carried out by PANAGENE Inc. (Daejeon, KR). The PNAs were dissolved in 100 μl of UltraPure™ DNase/RNase-Free Distilled water (Invitrogen™, Thermo Fisher Scientific, Waltham, MA, USA) to create a 200 μM stock solution. Subsequently, the PNAs stock solution was aliquoted and stored at −20 °C. For experimental use, the PNAs were diluted to 10 and 15 μM using a solution with 12 mM CaCl_2_ in UltraPure distilled water[34].

### Microinjection

Microinjections were performed using the Eppendorf^®^ FemtoJet^®^ Express microinjector. The GD-1 glass capillary with filaments (NARISHIGE Group. Tokyo, JP) was used for microinjection. To prevent clogging, the capillaries were coated with Sigmacote^®^ (Sigma-Aldrich^®^, Burlington, MA, USA). Fifteen glass capillaries were placed in a 15 ml centrifuge tube, to which one milliliter of Sigmacote^®^ was added. The tube was securely covered with a lid and subjected to gentle inversion to coat the glass capillaries evenly. Following coating, the glass capillaries were air-dried in a fume hood for 14 d. Prior to injection, glass capillaries were carefully pulled into sharp needles at 60 °C using a Narishige PC-10 puller (NARISHIGE Group. Tokyo, JP).

For the microinjection procedure, second instar aphid nymphs were immobilized on a microscopy glass slide layered with a piece of Kimwipe tissue paper (Misumi Group Inc., Tokyo, JP) moistened with 70 % ethanol, facilitated by a fine paintbrush. Microinjection was performed using a stereomicroscope (Leica S8 APO; Leica Microsystems, Wetzlar, DE). The injection pressure (Pi) was set to 1500 hPa, the injection duration (T) was 4 s, and the constant pressure (Pc) ranged between 400 and 700 hPa. The volume of PNAs injected into the aphid nymphs was estimated to be 0.2805 ± 0.0462 μL.

### The effect of peptide conjugated anti-*groEL* PNAs to *Buchnera* titer in the pea aphid

Aphid nymphs were injected with 15 μM of PNA_GroEL or PNA_mm. The injected aphid nymphs were kept in a plastic case containing broad bean seedlings. At 24 h post-injection, DNA was extracted from the injected aphid nymphs. A single injected aphid nymph was homogenized with a Nippi-sterilized Biomasher II (Funakoshi Co., Ltd., Tokyo, JP) in a tube containing 30 μl Buffer A (10 mM Tris-HCl, 1 mM EDTA, 25 mM NaCl in UltraPure distilled water) with 0.4 mg/ml Proteinase K (QIAGEN N.V., Venlo, NL). Homogenized samples were incubated at 37 °C for 1.5 h followed by 98 °C for 2 min. The extracted DNA samples were carefully stored in a freezer at −20 °C to ensure their preservation and integrity for subsequent analysis. The stored DNA samples were used to assess the effect of peptide-conjugated anti-*groEL* PNAs on *Buchnera* titer in pea aphids.

### Quantitative polymerase chain reaction (qPCR)

To quantify *Buchnera* titer in aphid nymphs treated with PNA_GroEL and PNA_mm, we performed the qPCR with KOD SYBR^®^ qPCR Mix (Toyobo Co., Ltd., Osaka, JP) using Roche LightCycler^®^ 96 real-time PCR instrument (Roche Applied Science, Penzberg, DE). The qPCR assay was conducted over 45 cycles, commencing with pre-incubation at 98 °C for 2 min followed by a 2-step amplification process. During this amplification, the DNA was denatured at 98 °C for 20 s, and then subjected to touchdown annealing from 66 to 57 °C, with a temperature decrement of 3 °C at each step, for 10 s. Subsequently, a 3-step amplification was applied, involving denaturation at 98 °C for 20 s, annealing at 66 °C for 10 s, and extension at 68 °C for 30 s. For the qPCR assay, we utilized the BuchDnaK_F1018 and BuchDnaK_R1142 primers, which specifically targeted the *dnaK* gene of *Buchnera*. Additionally, the rpL7_F and rpL7_R primers were used to amplify the *A. pisum rpL7* gene, which served as a normalization reference for data analysis (refer to Supplementary Table S1 for primer details)[35].

### The effect of peptide conjugated anti-*groEL* PNAs on *Buchnera groEL* expression

Aphid nymphs were subjected to injection with either 10 μM PNA_GroEL or PNA_mm. Following the injection, the treated aphid nymphs were housed in plastic cases, each containing a broad bean seedling. To investigate the effect of peptide-conjugated anti-*groEL* PNAs on *Buchnera groEL* expression, RNA was extracted from the injected aphid nymphs 24 h post-injection using the Takara Nucleospin RNA Plus XS RNA extraction kit (Takara Bio Inc., Shiga, JP). For RNA extraction, a single injected aphid nymph was homogenized with a Nippi-sterilized Biomasher II (Funakoshi Co., Ltd., Tokyo, JP) using Lysis Buffer I from the kit. RNA was extracted according to the manufacturer’s instructions.

### Reverse transcription-quantitative polymerase chain reaction (RT-qPCR)

The assessment of *Buchnera groEL* expression was carried out through the RT-qPCR assay, using the Takara One Step TB Green PrimeScript™ RT-PCR kit (Takara Bio Inc., Shiga, JP) and the Roche LightCycler® 96 real-time PCR instrument (Roche Applied Science, Penzberg, DE). The RT-qPCR procedure was initiated with a single round of reverse transcription at 42 °C for 5 min, followed by denaturation at 95 °C for 10 s. Subsequently, 30 cycles of PCR reactions were performed, involving denaturation at 95 °C for 5 s, followed by annealing and extension at 60 °C for 20 s. Finally, a 3-step melting curve analysis was applied, comprising denaturation at 95 °C for 10 s, annealing at 65 °C for 60 s, and extension at 97 °C for 1 s. For specific amplification of the *groEL* gene, we used the BuchGroEL_1_F and BuchGroEL_R primers, whereas the BuchDnaK_F1018 and BuchDnaK_R1142 primers were used to amplify the *Buchnera dnaK* gene, serving as a reference for normalization (see Supplementary Data Table S1 for primer details).

### Sample preparation for super-resolution laser scanning confocal microscopy imaging

Aphid nymphs were injected with a solution of 15 μM PNA_GroEL_TR or 12 mM CaCl2, and 24 h after microinjection, they were carefully dissected using fine needles (insect pins, Shiga Konchu Fukyusha, Japan) attached to wooden chopsticks in ice-cold 1x Phosphate buffered saline (PBS) (Gibco 10 x PBS pH 7.4, Thermo Fisher Scientific, MA, USA). Dissection was performed using a stereomicroscope. Subsequently, bacteriocytes, embryos, and other tissues were fixed with a 4 % paraformaldehyde phosphate buffer solution (Fujifilm Wako Pure Chemical Corp., Osaka, JP) for 30 min. After fixation, the tissues were washed thrice with ice-cold 1x PBS for 10 min each. The tissues were then stained with 1 μg/ml 4,6-diamidino-2-phenylindole (DAPI) in 1x PBS solution (Dojindo Laboratories, Kumamoto, JP) for two hours at room temperature.

Following staining, the tissues were washed thrice with ice-cold 1x PBS for 10 min each. Finally, the stained tissues were mounted on the glass slide with ProLong™ Gold antifade mountant (Invitrogen™, Thermo Fisher Scientific, Waltham, MA, USA). A #1.5 coverslip (Matsunami CS00802; Matsunami Glass Ind., Ltd., Osaka, JP) was used to cover the tissues on the slide. The microscope slides were mounted and cured at room temperature for 24 h before being sealed with clear nail polish.

To obtain single optical plane images, we used a Zeiss LSM 980 with Airyscan 2 in Airyscan SR mode. Visualization was performed by fluorescence and differential interference contrast (DIC) microscopy. Images were taken with a Plan-Apochromate 63x/1.40 oil immersion lens. Images were processed using Zen software in the default 2D super-resolution mode (Blue edition; Ver. 3.4.91). The imaging channels for Airyscan were set as follows: λ_ex_ = 561 nm (0.8 %, detector gain = 750 V) and detection wavelength = 380 – 735 nm for Texas Red; λ_ex_ = 473 nm (0.2 %, detector gain = 750 V) and detection wavelength = 380 – 735 nm for DAPI; λ_ex_ = 488 nm (0.2 %, detector gain = 350 V) and detection wavelength = 300 – 900 nm for DIC. Linear unmixing was performed using the Automatic Component Extraction technique, a built-in function of the Zen software, to remove autofluorescence.

### Data preparation for *Buchnera* count analysis using SymbiQuant

The dissection, fixation, DAPI staining, and mounting of bacteriocytes were performed following the previously described method. However, instead of using 1x PBS, we used a 0.2 % Triton® X-100 (Sigma-Aldrich®, Burlington, MA, USA) in 1x PBS (Ptx) solution to facilitate the dissociation of the bacteriome into individual bacteriocytes. Subsequently, single optical plane images were acquired using an Olympus FluoView FV1000 confocal microscope, selecting the plane in which the nucleus of the bacteriocyte exhibited the widest dimension. Visualization was performed using fluorescence and differential interference contrast microscopy. For DAPI imaging, the imaging channel was set at λex = 405 nm and λem = 430 – 455 nm, while differential interference contrast microscopy was conducted using a 559 nm laser. Images were acquired using a 100x UPlanApo, 1.45NA/0.13WD oil immersion lens. After acquisition, images were processed using FV10-ASW (Ver. 3.0) software. The DAPI channel of each image was exported at a resolution of 1024 x 1024 pixels. tiff format. To measure the area covered by *Buchnera* cells, we used the ImageJ Wand Tool (Ver. 1.52q) [36]. Quantification of the *Buchnera* count was performed using SymbiQuant software in Google Colab (https://github.com/EBJames/SymbiQuant), following the authors’ instructions[23]. The *Buchnera* count for each bacteriocyte was normalized by the bacteriocyte area (µm^2^) measured with ImageJ.

### The effect of peptide conjugated anti-*groEL* PNAs on *Buchnera pyrG* expression

We performed RT-qPCR to check the expression of *Buchnera pyrG* in PNA_GroEL-and PNA_ mm-treated aphid nymphs. The RT-qPCR protocol has been previously described. For gene amplification, we used BuchPyrG_F and BuchPyrG_R primers, which specifically target the *pyrG* gene of *Buchnera*. To normalize the data, BuchDnaK_F1018 and BuchDnaK_R1142 primers were used to amplify *Buchnera dnaK*. Moreover, we analyzed the expression of *Buchnera rrs* and *rpoA* in the PNA_GroEL-and PNA_ mm-treated groups using the same RT-qPCR method. Primer details are provided in Supplementary Data Table S1.

### Statistical analysis

qPCR data analysis was performed using a one-tailed Student’s t-test to compare the effects of PNA_GroEL and PNA_mm treatments on *Buchnera* titers in aphid nymphs. Specifically, one-tailed Student’s t-tests were used to examine the differences in *Buchnera groEL*, *rrs*, and *rpoA* expression between the PNA_GroEL-and PNA_ mm-treated groups. In contrast, a two-tailed Mann-Whitney U test was used for *pyrG* expression analysis. The *Buchnera* count obtained from SymbiQuant was normalized with the bacteriocyte area followed by log transforming the normalized count (Log_10_(*Buchnera* count/ µm^2^)) before conducting a one-tailed Welch Student’s t-test to compare the results between the two groups. All data analyses were performed using the R version 4.2.2 and relevant packages, including “devtools,” “DescTools,” and “dplyr”[37]. For data visualization, boxplots were generated using the “ggpubr” and “ggsignif” packages in R. Additionally, survival analysis was carried out using various packages such as “survival,” “lubridate,” “gtsummary,” “tidycmprsk,” “condSURV,” and “survminer.” Kaplan-Meier curves were plotted using the “ggsurvfit” package. Furthermore, a log-rank test was used to compare the survival times of PNA_GroEL-and PNA_ mm-treated aphid nymphs.

### Rights and permissions for research involving plants

This study did not require special right or permissions for plant material use. The seeds of broad bean plants (*Vicia faba* L.) used for culturing aphids in this study were not from wild plants, but were obtained commercially. These seeds are intended for human consumption or as feed for domestic animals, such as race pigeons. This plant is not categorized as an endangered species. All methods and analyses were carried out in accordance with relevant institutional, national, and international regulations and guidelines.

## Data availability

All relevant data are within the manuscript and its Supporting Information. Raw data used in this study are available from the corresponding author upon request.

## Acknowledgements

We express our gratitude to members of the Laboratory of Evolutionary Genomics for their invaluable support and constructive advice. We appreciate the technical advice provided by Prof. Kazuya Kobayashi of Hirosaki University and Dr. Kiyono Sekii of Keio University. We thank the Optics and Imaging Facility, NIBB Trans-Scale Biology Center, for their technical support.

## Author contributions

K.T. and S. S. designed the study; K. T. performed the experiments and analyzed the data; K.T. and S. S. wrote the manuscript and contributed to its revision.

## Funding

This work was funded by the Japan Society for the Promotion of Science (JSPS) KAKENHI Grants-in-Aid for Scientific Research (A) (No. JP20H00478) to S. S. and MEXT Scholarship to K. T.

## Competing interests

There are no conflicts of interest to declare.

## Notes

### Competing Interest Statement

The authors have declared no competing interest.

