## Supplementary for "Synthetic Peptide Nucleic Acids in Action: *In Vivo* Interference of the *groEL* Gene from Pea Aphid Endosymbiont *Buchnera aphidicola*"

### Supplementary data

Table S1: List of primers used in qPCR and RT-qPCR analyses.

| Primer | Sequence | Primer length (bp) | Amplicon length (bp) | Target gene | Reference |
| --- | --- | --- | --- | --- | --- |
| BuchGroEL_1_F | AAACTATCAGGCGGTGTTGC | 20 | 226 | *groEL* |  |
| BuchGroEL_R | CACGCAAAGCAACTCGAATA | 20 |  |  |  |
| rpL7_F | GCGCGCCGAGGCTTAT | 16 | 81 | *rpl7* |  |
| rpL7_R | CCGGATTTCTTTGCATTTCTTG | 22 |  |  | [*] |
| BuchPyrG_F | AATAGGCGGAACTGTTGGTG | 20 | 248 | *pyrG* |  |
| BuchPyrG_R | TTTTTCGCTCGTGAAGAGGT | 20 |  |  |  |
| BuchDnaK_F1018 | GTTGGTGGTCAAACTAGAATGCCT | 24 | 125 | *dnaK* |  |
| BuchDnaK_R1142 | ACTCCTCCCTGTACTGCAGC | 20 |  |  |  |
| Buch_rpoA_F | CAGGATGTGCGGTAACTGAA | 20 | 185 | *rpoA* |  |
| Buch_rpoA_R | TGCGGCAGTAATAGAACCAA | 20 |  |  |  |
| Buch_rrs_F274 | AGGATAACCAGCCACACTGG | 20 | 115 | 16S rRNA gene |  |
| Buch_rrs_R388 | TCTTCATACACGCGGCATAG | 20 |  |  |  |

[*]	Nakabachi, A., Shigenobu, S., Sakazume, N., Shiraki, T., Hayashizaki, Y., Carninci, P., ... & Fukatsu, T. (2005). Transcriptome analysis of the aphid bacteriocyte, the symbiotic host cell that harbors an endocellular mutualistic bacterium, *Buchnera*. *Proceedings of the National Academy of Sciences*, *102*(15), 5477-5482.

Table S2: Sequence matches of anti-*groEL* PNAs in *Buchnera aphidicola* str. APS genome.

| Gene Name | CDS start position | CDS end position | Strand | Query start position | Query end position | Query strand | Comment | Extra Comment | Query sequence |  |
| --- | --- | --- | --- | --- | --- | --- | --- | --- | --- | --- |
| WP_009873980.1 (*groEL*) | 18715 | 20361 | + | NC_002528.1 | 18710 | 18719 | -1 | reverse complement match | Translational Start Site Included | GTCAAATGGC |
| WP_010896104.1 (*pyrG*) | 451384 | 453021 | + | NC_002528.1 | 451856 | 451865 | -1 | reverse complement match |  | GTCAAATGGC |

Figure S1

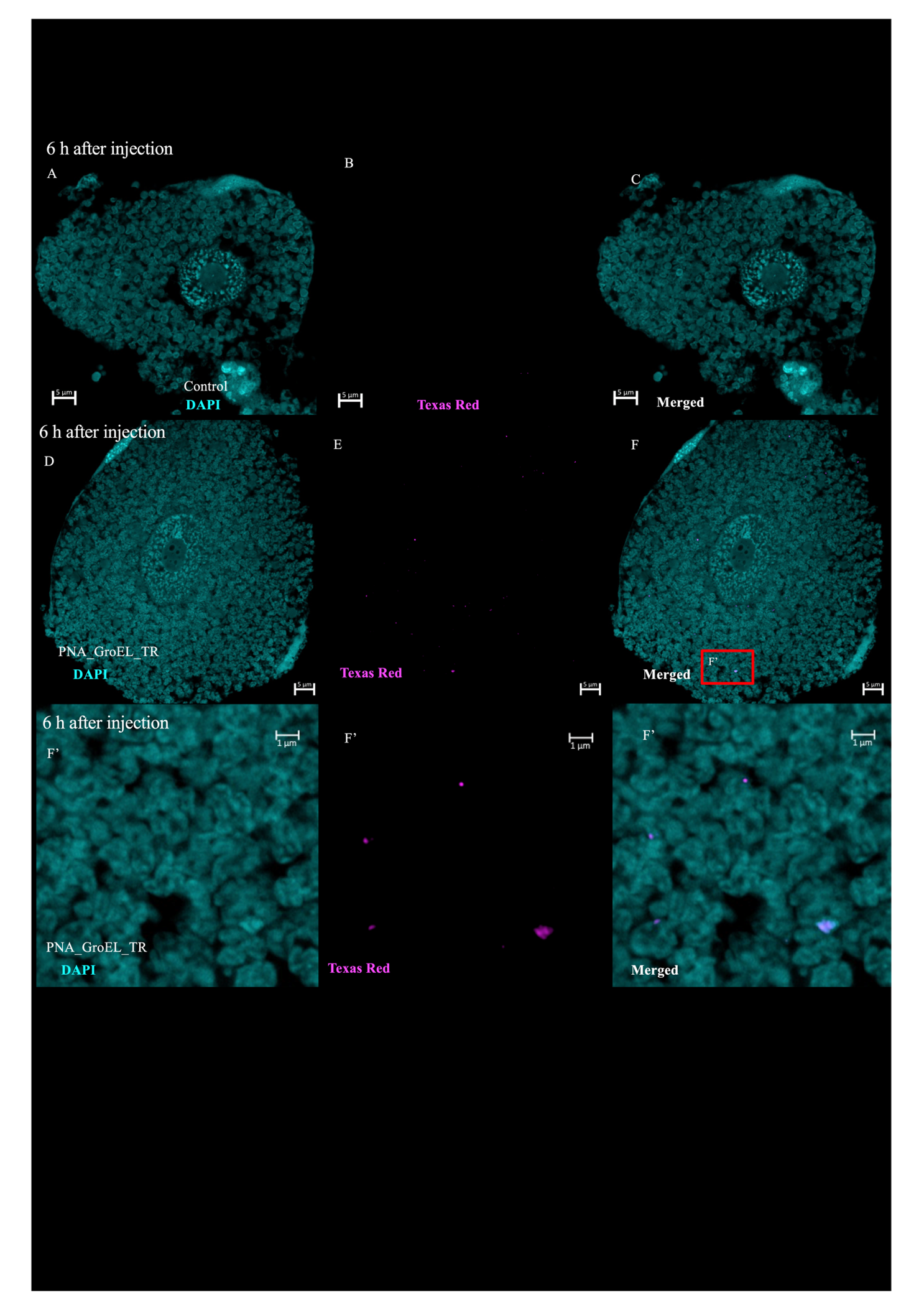

Figure S1: Super-resolution imaging of aphid bacteriocytes and *Buchnera* cells showed the signal of Texas Red which indicates the successful penetrated of PNA_GroEL (10 μM) into bacteriocytes 6 h after injection (D-F). CaCl_2_ solution (12 mM) injected samples were used as the control in the experiment (A-C). Zoom-in image clearly showed the Texas Red signal and distorted *Buchnera* in the bacteriocytes of PNA_GroEL_TR treated sample (F’).

Figure S2

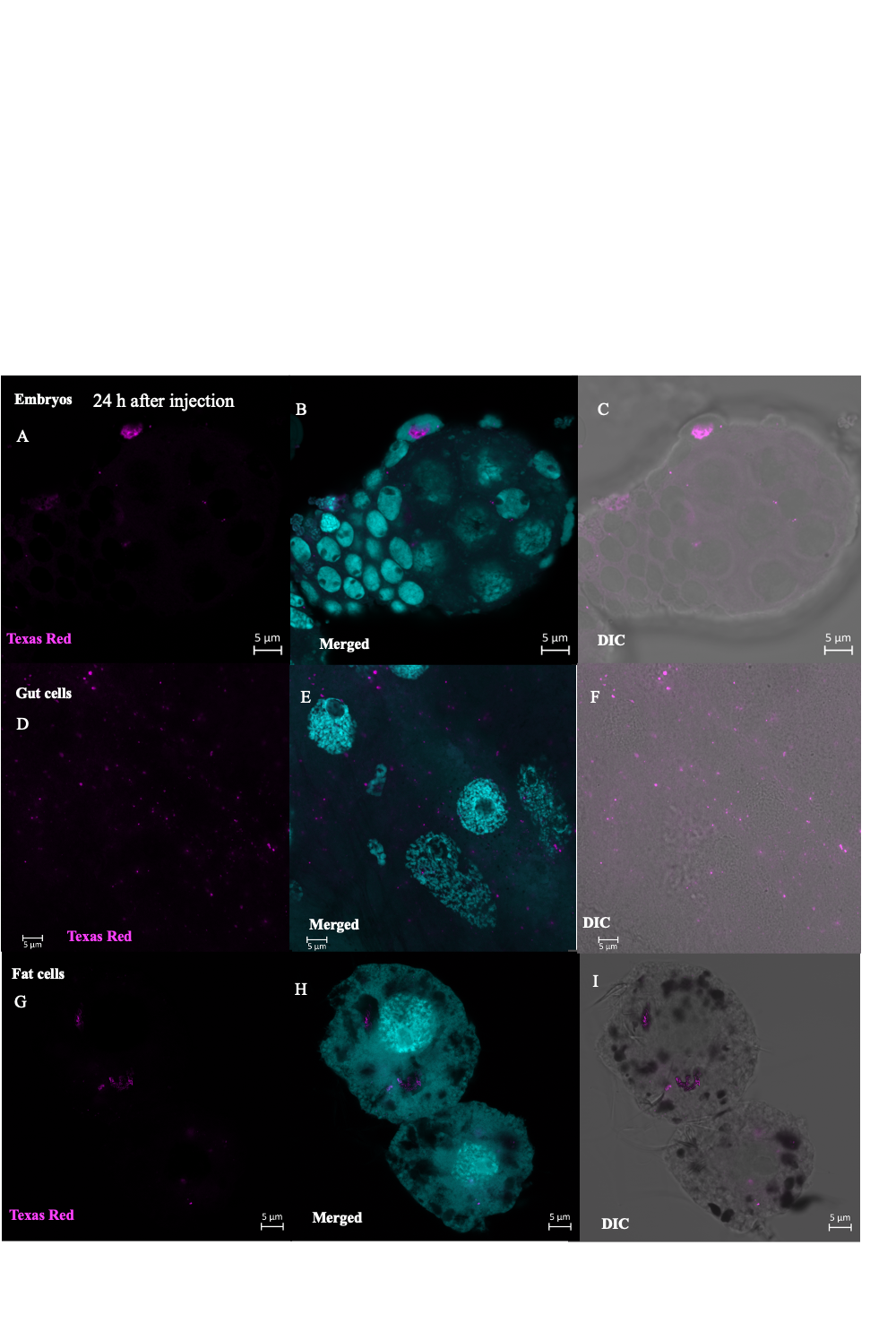

Figure S2: Super-resolution imaging of aphid bacteriocytes and *Buchnera* cells showed that the signal of Texas Red was detected in other aphid tissue such as fat cells, gut cells and embryos, in PNA_GroEL_TR (10 μM) treated samples (A-I). The images were taken using the aphid nymphs dissected after 24 h of PNA_GroEL_TR injection.

Figure S3

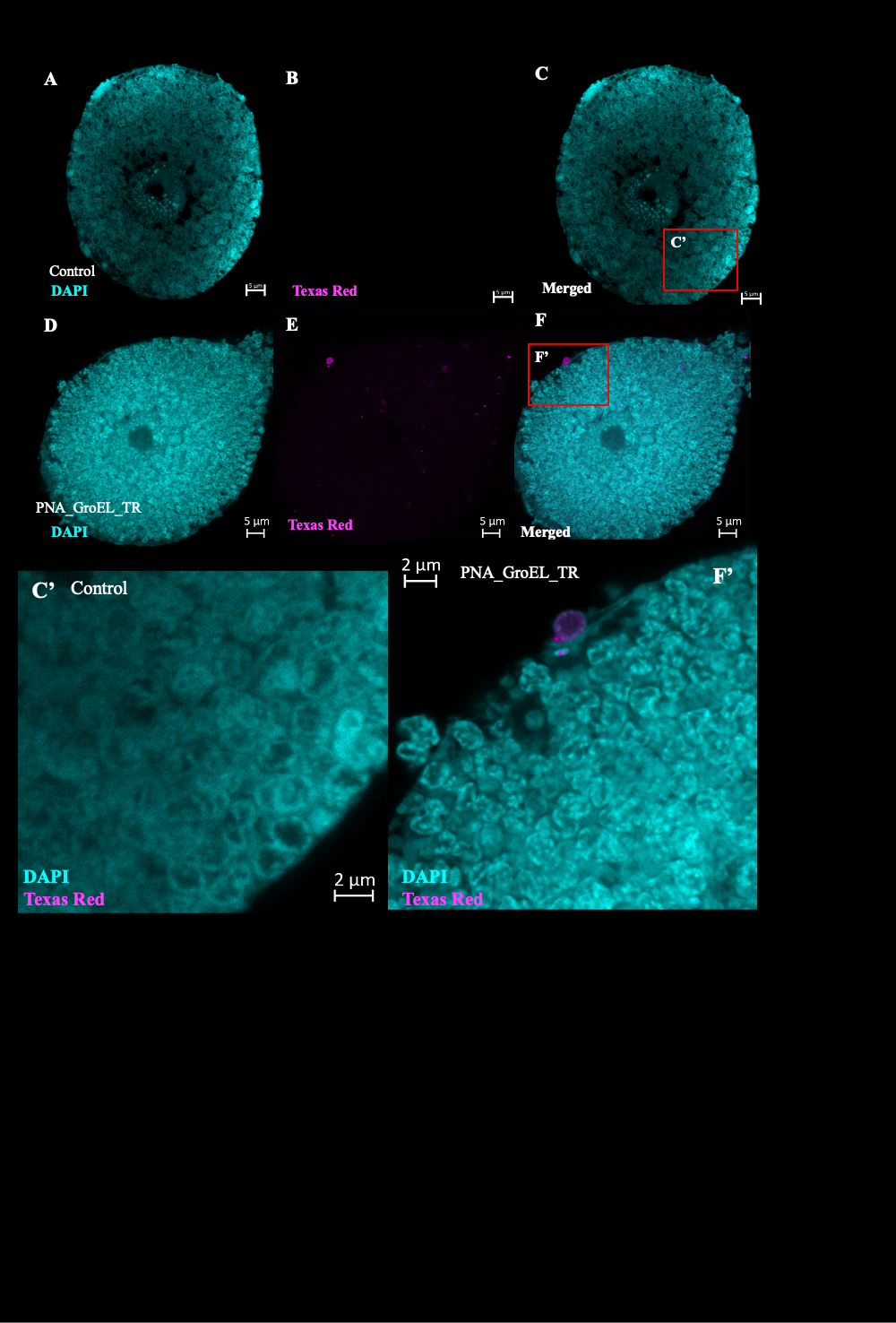

Figure S3: Super-resolution imaging of aphid bacteriocytes and *Buchnera* cells. Aphid nymphs were injected with CaCl_2_ solution (12 mM) or PNA_GroEL_TR (10 μM in 12 mM CaCl_2_ solution). Injected aphid nymphs were dissected after 24 h. Dissected bacteriocytes were fixed and stained with DAPI solution to mark the nuclei of bacteriocytes and chromosome of *Buchnera*. Aphid nymphs injected with CaCl_2_ solution injected was used as the control (A-C). Detection of the Texas Red signal in bacteriocytes and *Buchnera* cells indicated the successful penetration of PNAs into the cells (D-F). Zoom-in images showed that the *Buchnera* cells of CaCl_2_ treated aphid nymphs were mostly having a round shape (C’) while distorted *Buchnera* cells were found in PNA_GroEL_TR treated sample (F’).

### Figure S4 Expression of *Buchnera rrs* and *rpoA* genes were not inhibited by the treatment of peptide-conjugated anti-*groEL* PNAs

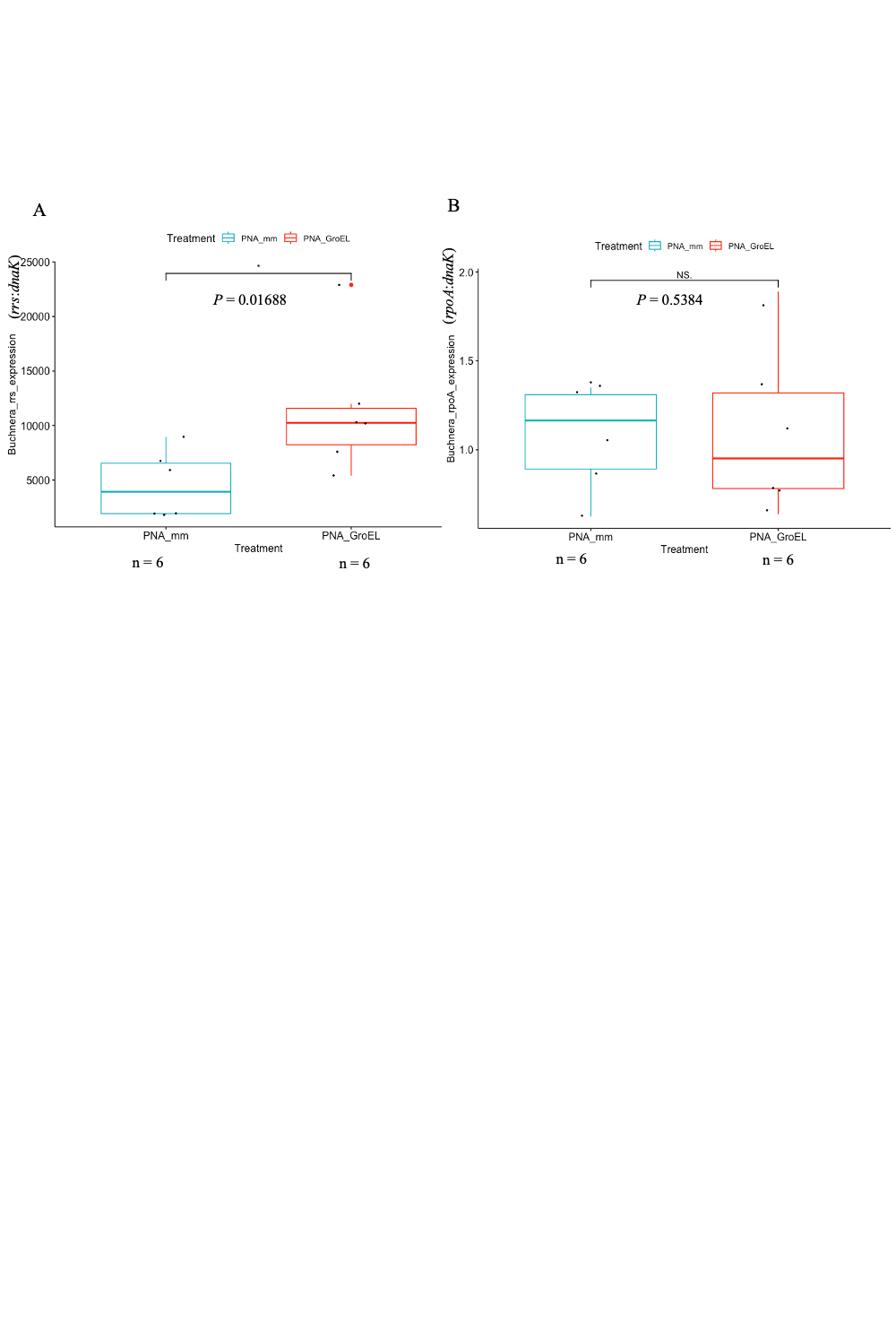

Figure S4: Comparison of *Buchnera* *rrs* and *rpoA* gene expression in PNAs injected aphid nymph. Second instar aphid nymphs were injected with 10 μM peptide-conjugated antisense *groEL* PNA (PNA_GroEL; red boxplot) or control PNAs (PNA_mm; blue boxplot). A significant higher expression of *rrs* gene was found in the aphid nymphs treated with PNA_GroEL than those treated with PNA_mm (One-tailed Student’s t-test, *p* = 0.01688 < 0.05) while the difference in *Buchnera rpoA* gene expression between aphid nymphs treated with PNA_GroEL and PNA_mm was not significant (One-tailed Student’s t-test, *p* = 0.5384 > 0.05). The *p* value is shown between the two boxplots. Each black dot indicates a single PNAs treated aphid nymph.

We compared the expression of other *Buchnera* genes, i.e. *rrs* and *rpoA*, to determine whether the treatment of peptide-conjugated anti-*groEL* PNAs has inhibited the expression of untargeted genes. A comparison of *Buchnera* *rrs* gene expression between PNA_GroEL and PNA_mm treated aphid nymphs was carried out by RT-qPCR (Figure S4A). Meanwhile, the comparison of *Buchnera rpoA* gene expression between the two treatments is shown in Figure S4B. The expression of *rrs* and *rpoA* genes were normalised by *Buchnera dnaK* gene expression. *Buchnera rrs* gene expression in aphid nymphs treated with 10 μM PNA_GroEL (*M* = 11400, *SD* = 6093.541) was significantly higher than those treated with the same concentration of PNA_mm (*M* = 4548.333, *SD* = 3078.126) (*t* (10) = 2.4584, *p* = 0.01688). On the other hand, no significant difference was detected between PNA_GroEL and PNA_mm in terms of the expression of the *rpoA* gene (*M* = 1.10, *SD* = 0.472 for PNA_GroEL; *M* = 1.08, *SD* = 0.295 for PNA_GroEL) (*t* (10) = 0.098979, *p* = 0.5384).

### Figure S5

### Treatment of peptide-conjugated anti-*groEL* PNAs did not influence the probability of aphid nymphs’ survival

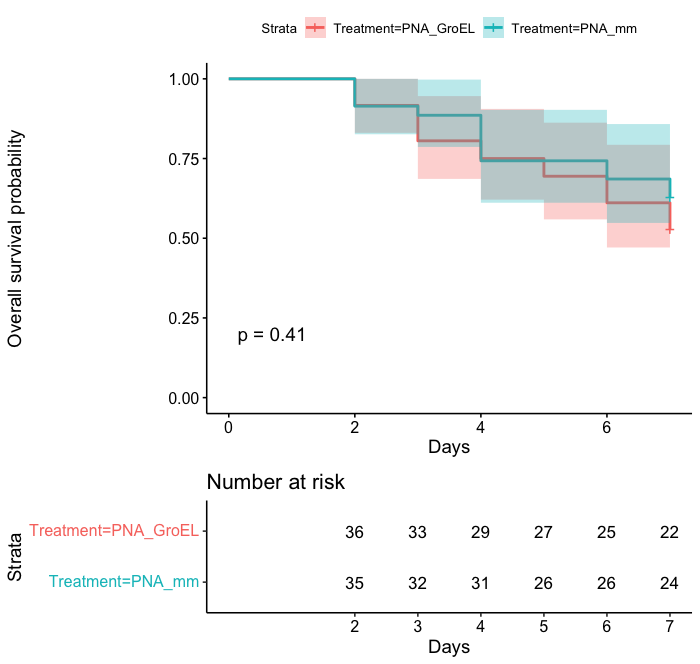

Figure S5: Kaplan-Meier survival curves of aphid nymphs treated with PNA_GroEL and PNA_mm. Second instar aphid nymphs were injected with 10 μM peptide-conjugated antisense *groEL* PNA (PNA_GroEL; red boxplot) or control PNAs (PNA_mm; blue boxplot), 36 and 35 aphid nymphs respectively. There is no significant difference in survival between PNA_GroEL and PNA_mm treated aphid nymphs (*p =* 0.4 > 0.05, log-rank test).

We performed a log-rank test to determine whether there is a significant difference in survival times between PNA_GroEL and PNA_mm treated aphid nymphs. Kaplan-Meier plots were showed in Figure S5. Survival of aphid nymphs was observed for seven days after being treated with PNA_GroEL or PNA_mm. We found that 61.1 % of PNA_GroEL-treated aphids and 68.6 % of PNA_mm-treated aphid nymphs survived more than seven days. The log-rank test result showed no significant difference between the survival of PNA_GroEL and PNA_mm treated aphid nymphs (*p =* 0.41).

### Access link of Google Colab notebook for finding the sequence match of PNAs:

https://colab.research.google.com/drive/1J8flkB8qwORqKCCLrDnBlOxKlVGtP1uF?usp=sharing
